# Local axonal conduction delays underlie precise timing of a neural sequence

**DOI:** 10.1101/864231

**Authors:** Robert Egger, Yevhen Tupikov, Kalman A. Katlowitz, Sam E. Benezra, Michel A. Picardo, Felix Moll, Jörgen Kornfeld, Dezhe Z. Jin, Michael A. Long

**Author notes:** To whom correspondence should be addressed: Michael A. Long. Authors contributed equally (Robert Egger, Yevhen Tupikov).

## Abstract

Sequential activation of neurons has been observed during various behavioral and cognitive processes and is thought to play a critical role in their generation. Here, we studied a circuit in the songbird forebrain that drives the performance of adult courtship song. In this region, known as HVC, neurons are sequentially active with millisecond precision in relation to behavior. Using large-scale network models, we found that HVC sequences could only be accurately produced if sequentially active neurons were linked with long and heterogeneous axonal conduction delays. Although such latencies are often thought to be negligible in local microcircuits, we empirically determined that HVC interconnections were surprisingly slow, generating delays up to 22 ms. An analysis of anatomical reconstructions suggests that similar processes may also occur in rat neocortex, supporting the notion that axonal conduction delays can sculpt the dynamical repertoire of a range of local circuits.

## INTRODUCTION

Sequential neural activity in local brain areas is thought to play a critical role in behaviors such as motor control (Luczak et al., 2015; Mauk and Buonomano, 2004; Peters et al., 2014; Prut et al., 1998), navigation (Foster and Wilson, 2007; Pastalkova et al., 2008), and decision-making (Mello et al., 2015; Schmitt et al., 2017). A variety of mechanisms have been proposed to underlie the generation of neural sequences (Diesmann et al., 1999; Fiete et al., 2010; Goldman, 2009; Hahnloser et al., 2002; Kleinfeld and Sompolinsky, 1988; Laje and Buonomano, 2013; Rajan et al., 2016), but experimental tests of these network models have been stymied by the scarcity of data sets that relate behavior, network function and circuit structure. The zebra finch is an advantageous model organism for studying the network basis of neural sequence generation. Each adult male zebra finch produces a courtship song that is nearly identical from one rendition to the next, consisting of ~3-7 discrete vocal elements known as ‘syllables’ (Figure 1A). Many lines of evidence have suggested that neural activity controlling the moment-to-moment timing of song production is localized to a single brain region, called HVC (proper name) and is driven by premotor neurons in that region (Figure 1B) (Hahnloser et al., 2002; Long and Fee, 2008; Nottebohm et al., 1976; Vu et al., 1994). HVC premotor neurons produce high-frequency bursts of action potentials (~4-5 spikes/burst, ~10 ms duration) at a single moment during the song (Hahnloser et al., 2002) with millisecond precision across song renditions (Figure 1C). At the network level bursts form a sustained sequence spanning the duration of song syllables (Figure 1D and 1E).

**Figure 1.**
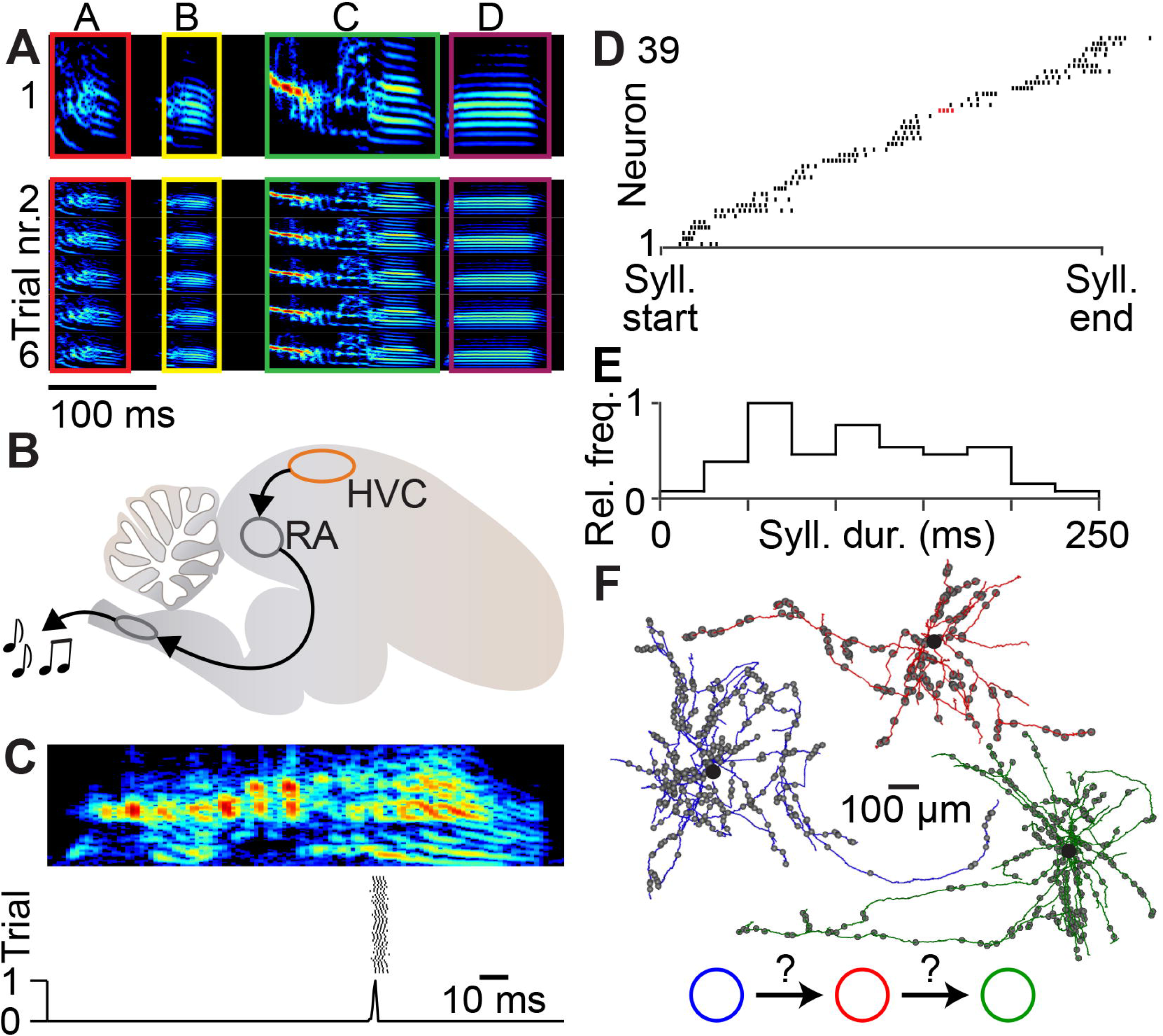
A precise neural sequence underlies zebra finch song production. (A) Spectrogram of an example song motif consisting of four discrete syllables (top) and five different repetitions of the same song motif (bottom). (B) An illustration of the zebra finch brain showing nucleus HVC and its downstream target along the song production pathway, the robust nucleus of the arcopallium (RA). RA sends projections to brainstem motoneurons involved in producing vocalizations. (C) Spectrogram of an individual syllable (top) and spike raster plots of an HVC premotor neuron during different song renditions (middle) (Lynch et al., 2016). Bottom: relative frequency of burst onset times across trials. (D) Representative spike trains from different neurons aligned relative to on-offset of the syllable during which they are active. Red: example spike train from (C). (E) Distribution of syllable durations (58 syllables from 14 birds). (F) Example reconstruction of the local (i.e., within HVC) axon collaterals of three HVC premotor neurons. Black circle: soma location; grey spheres: modeled location of synapses onto other HVC premotor neurons along the axon.

What are the synaptic interactions enabling HVC sequence generation? Previous anatomical (Kornfeld et al., 2017) and electrophysiological (Kosche et al., 2015; Mooney and Prather, 2005) studies had demonstrated direct excitatory connections between premotor neurons (Figure 1F), but the sufficiency of these connections for propagating activity from earlier to later steps in a sequence remains controversial (Cannon et al., 2015; Galvis et al., 2018; Gibb et al., 2009; Hamaguchi et al., 2016; Jin et al., 2007; Long et al., 2010; Pehlevan et al., 2018). For instance, activity may be driven through single strong connections (Lorteije et al., 2009) or through convergent inputs from several presynaptic partners (Bruno and Sakmann, 2006). Two primary lines of evidence support the latter possibility. Local excitatory synaptic strength, as estimated by active zone size in our previous ultrastructural work (Kornfeld et al., 2017), is not extraordinarily large, which does not support the idea of single presynaptic partners. Consistent with this view, unitary synaptic potentials measured with paired intracellular recordings (~2 mV) are considerably smaller than the depolarization observed during singing (~10 mV), suggesting that many presynaptic elements are involved in the generation of spiking events (Kosche et al., 2015; Long et al., 2010; Mooney and Prather, 2005). Although these presynaptic partners are likely to be other premotor neurons within HVC, their exact identity (i.e., spatial location, specific timing) remains unknown. Therefore, in order to take a step towards resolving the present conflicting theories about sequence generation in HVC, it is necessary to understand the nature of this functional convergence.

To address this issue, we used a network model constrained by experimental measurements of HVC premotor neuron properties and population activity. We found that the spatiotemporal organization of the HVC sequence during singing is best matched by a neural network architecture in which neurons are connected by local axon collaterals with long conduction delays – matching those observed in HVC. Because of these heterogeneous conduction delays, sequential activity propagates via convergent input from presynaptic neurons active at different times and, as a result, activity is ‘polychronous’ (Izhikevich, 2006) - generating continuous time-locked spiking patterns without synchrony. To assess the significance of local axonal delays in other brain areas, we estimated conduction delays along axons of layer 4 (L4) neocortical neurons (Narayanan et al., 2015), and find that the differences in axonal arbor size and conduction velocity compensate and yield delays similar to those observed in zebra finch HVC. Hence, axonal conduction delays may play a key role in shaping network activity within local brain circuits.

## RESULTS

### Synfire chain models do not explain HVC population data

An effective mechanism for generating postsynaptic convergence is the synchronous activation of presynaptic neurons (Figure 2A) (Bruno, 2011; Bruno and Sakmann, 2006). Feedforward networks based on synchronous neuronal activation, known as ‘synfire chains’, have long been suggested to underlie sequence generation (Abeles, 1991; Amari, 1972). This network architecture has previously been proposed to explain HVC sequences (Fee et al., 2004; Fiete et al., 2010; Jin et al., 2007; Long et al., 2010; Pehlevan et al., 2018), and has the benefit of capturing the precision of individual neurons and the stability of network sequences (Jin et al., 2007; Long et al., 2010). We generated a synfire chain network model of 20,000 active HVC premotor neurons (Long et al., 2010), where HVC premotor neurons were triggered by synchronously active presynaptic ensembles (see Methods), as reflected in the fine structure of the network output (Figure 2B, i.e., discrete groups of synchronously active neurons with ~5-6 ms intervals between burst onset times). As before, this model produced a sustained network sequence covering syllable-length timescales.

**Figure 2.**
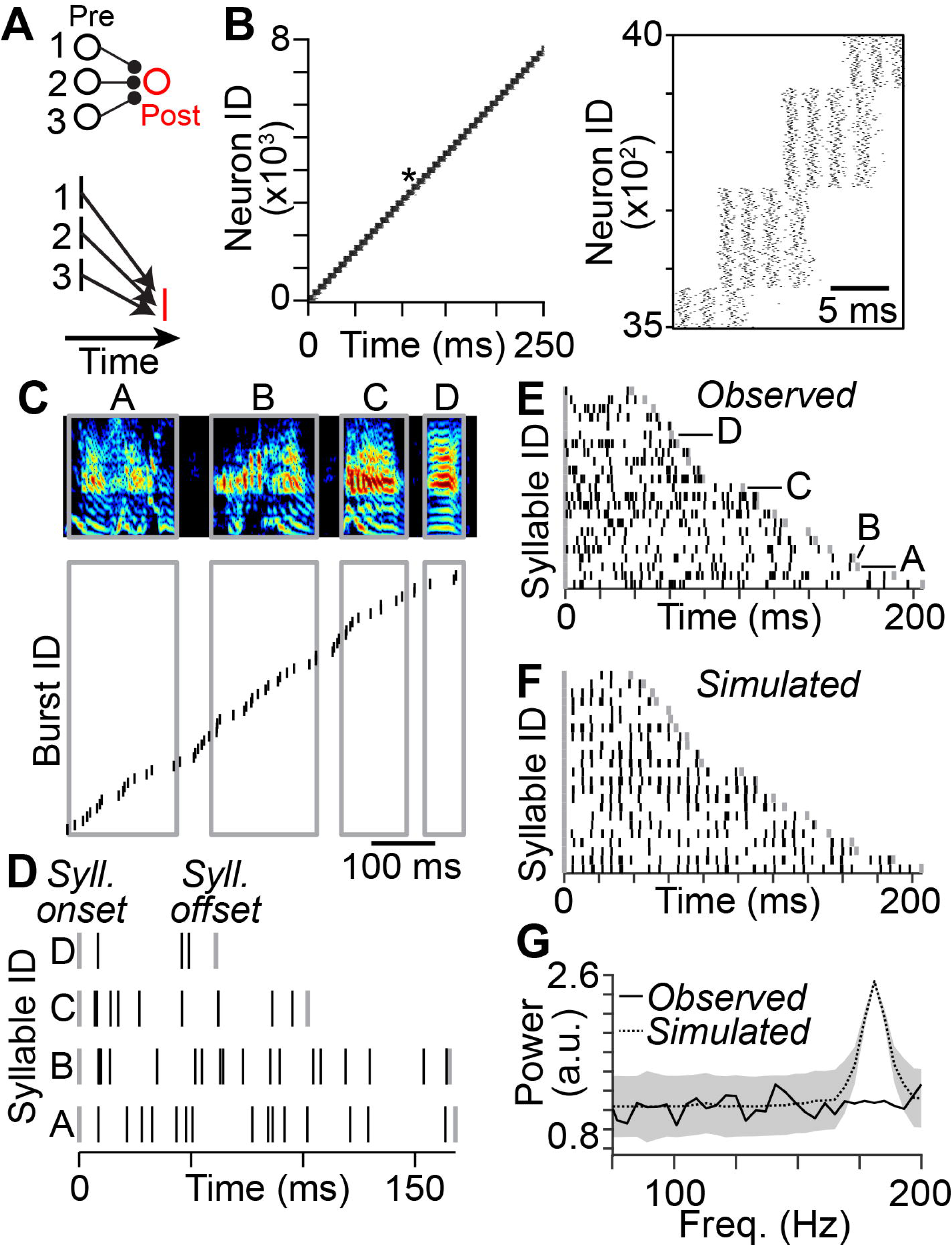
Synfire chain model of HVC premotor neurons predicts synchronized network sequences not observed *in vivo*. (A) Synchronously active groups of neurons form convergent synaptic connections onto the same postsynaptic neuron, resulting in a sustained sequence at the population level. (B) Spike raster plot of sequential activity in a synfire chain model of HVC premotor neurons with numbers of synaptic connections constrained by anatomical measurements. Spikes from 10% of all active neurons are shown. Inset: Magnified view of spike raster plot highlighted (*), revealing synchronously active groups of neurons. (C) Top: Spectrogram of song consisting of four syllables. Bottom: Burst onset times of HVC projection neuron activity recorded during song. Grey: syllables. (D) Burst onset times in (C) relative to on- and offset of the syllables during which they occur. (E) Burst onset times of HVC projection neurons relative to syllable on- and offset in 23 syllables from 5 birds (Lynch et al., 2016). Syllables from (D) highlighted. (F) Burst onset times predicted by the synfire chain model by matching duration and number of observed burst times for each syllable. (G) Power spectrum of burst onset times calculated from experimental observations and the synfire chain model. Shaded area: ±3 SD (bootstrap).

Until recently, such network models were difficult to test empirically; the number of recordings performed in HVC during song production were limited (Amador et al., 2013; Hahnloser et al., 2002). However, following recent improvements in recording technology, we can now track the activity of large cell populations (Lynch et al., 2016; Okubo et al., 2015; Picardo et al., 2016) to test model predictions against the data. Therefore, we looked for this pattern of synchronized burst onset times in HVC by examining a recently reported data set of 286 projection neurons in 5 birds measured using extracellular recordings during singing (Figure 2C and 2D) (Lynch et al., 2016). We compared these measurements with predictions from the synfire chain model (Figure 2E and 2F) and found that the timing of activity was qualitatively different: the synfire model predicted that activity would be clustered at distinct timepoints, while the recorded data appeared to lack such temporal structure. To obtain a quantitative comparison of these differences, we computed the power spectrum of burst onset times. The synfire chain model predicted a peak in the power spectrum at ~180 Hz, corresponding to the ~5-6 ms interval between synchronous groups of neurons (Figure 2B), while the recorded data exhibited a flat power spectrum, consistent with a more uniform distribution of burst times (Figure 2G). Therefore, HVC activity does not appear to be restricted to synchronous groups, suggesting that the network connectivity is not organized as a synfire chain.

### Temporal structure of polychronous network sequences is affected by conduction delays

In an alternative model, presynaptic neurons are active across a range of different times, and heterogeneous network delays are sufficient to allow spikes from multiple neurons to arrive simultaneously at a postsynaptic target (Figure 3A) (Izhikevich, 2006). Previous theoretical work based on conduction delays measured between different brain areas (Swadlow, 1985, 1994) demonstrated that such a ‘polychronous’ network architecture can generate time-locked spiking patterns without synchrony (Bienenstock, 1996; Izhikevich, 2006), but it is unclear if delays in local circuits are sufficient to generate sustained sequences in this context. To estimate these delays, we needed a measure for axonal pathlength as well as conduction velocity. Axonal pathlengths to each synapse were determined using previous observations from our laboratory; the relative position and abundance of synapses were measured using electron microscopy (Kornfeld et al., 2017) and the total axonal extent for each neuron was obtained from 22 complete light microscopic reconstructions of local axonal collaterals (Figure 3B) (Benezra et al., 2018). Our first estimate for conduction velocity was 0.3 mm/ms, a value reported for local unmyelinated axons in mammalian neocortex (Helmstaedter et al., 2008; Hirsch and Gilbert, 1991; Shu et al., 2007). Using these parameters, we developed a procedure to generate a feedforward polychronous network of HVC premotor neurons for a given distribution of axonal conduction delays (Figure S1). Because measured pathlength distances follow a log-normal distribution (Buzsaki and Mizuseki, 2014), we use this shape as our first estimate of these delays (Figure S1F). We find that the resulting network produces a reliable sequence in which neurons are active in a continuous fashion with burst onset times spread throughout (Figure 3C), thus matching our empirical observations (Figures 2E, 3D and 3E), in contrast to the discrete synchronous activity of the synfire chain model (e.g., Figure 2F and 2G).

**Figure 3.**
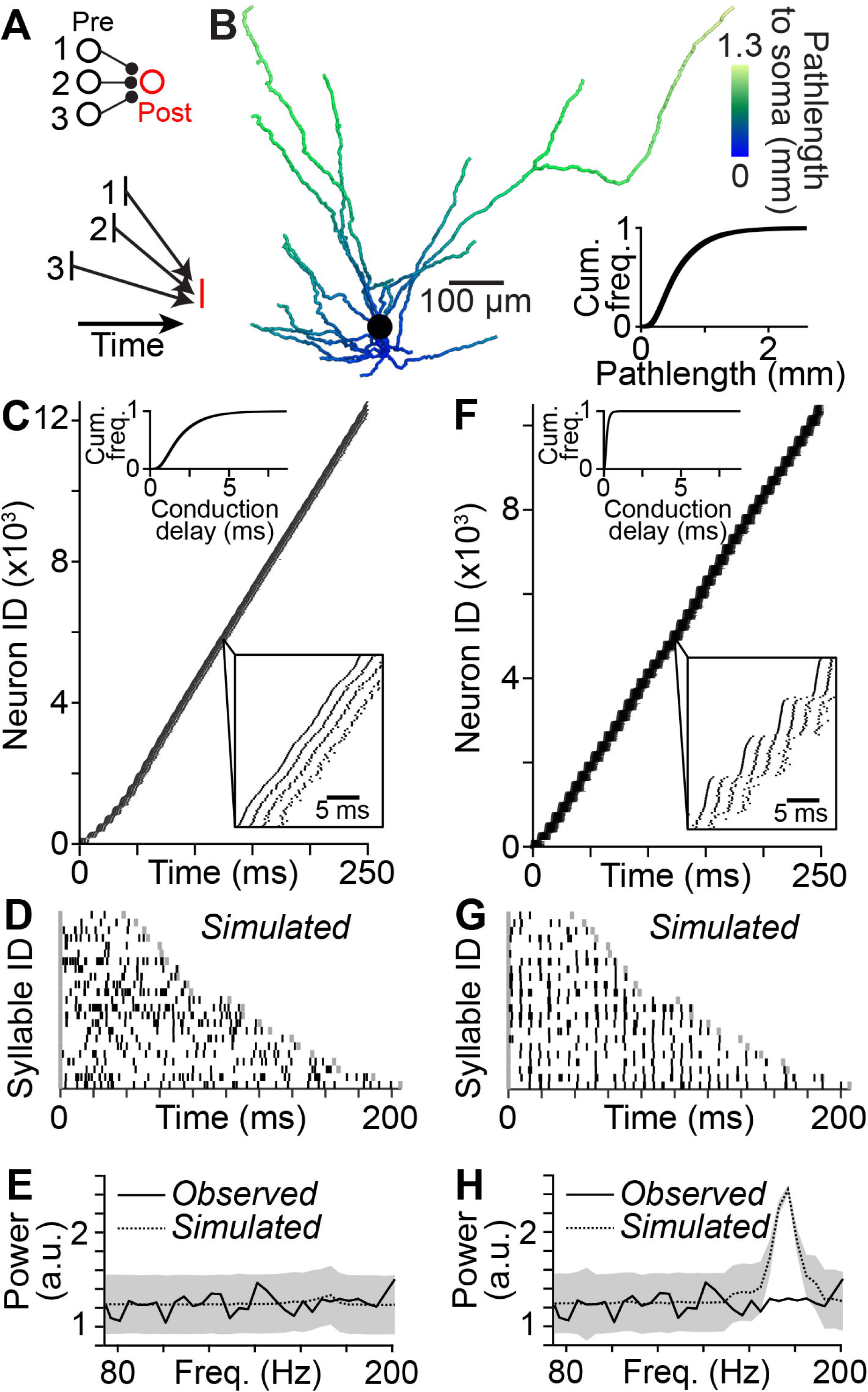
Heterogeneous axonal conduction delays enable continuous network sequences in polychronous network models. (A) The activity of neurons forming convergent synaptic connections onto the same postsynaptic neuron does not have to be synchronized. Instead, if differences in spike times are compensated by suitable delays, these spikes arrive synchronously at the postsynaptic neuron, resulting in a network sequence without synchrony. (B) Pathlength to soma measured along the reconstruction of local axon collaterals of an HVC premotor neuron. Inset: Pathlength distribution measured for 22 HVC premotor neuron axons. (C) A polychronous network model of HVC premotor neurons connected by synapses with conduction delays (top inset) based on axonal pathlengths and a conduction velocity of 0.3 mm/ms generates a sustained sequence of bursting activity. Bottom inset: Burst onset times are distributed continuously throughout the sequence. Spikes from 10% of all active neurons are shown. (D) Burst onset times predicted by the polychronous network model by matching duration and number of observed burst times for each syllable. (E) Power spectrum of burst onset times calculated from the observed burst onset times and the polychronous network model. Shaded area: ±3 SD (bootstrap). (F) A polychronous network model of HVC with ten times smaller conduction delays compared to the delays in C (top inset) results in a sequence of bursting activity of HVC premotor neurons in which bursts occur in synchronous groups of neurons (bottom inset). (G, H) As in D and E for the above model in F.

How important are axonal conduction delays in the function of the polychronous network? To examine this question, we artificially increased the conduction velocity of local axons by an order of magnitude within our network model. When we made the axonal delays ten times shorter (Figure 3F), we found that the resulting network sequence consisted of groups of neurons bursting near-synchronously (Figure 3F–3H), as in the synfire chain model. This result suggests that the nature of sequential activity can qualitatively shift based on the distribution of axonal conduction delays. To explore this issue more thoroughly, we created a range of models in which the means and standard deviations (SD) of axonal delays were varied parametrically over an order of magnitude (Figures 4A, 4B, and S1F), encompassing the examples discussed above (Figure 3). For each combination of mean and SD, we generated a polychronous network model and simulated network activity. We then compared our model with experimental observations by determining whether the degree of network synchrony in each model – as determined by the power spectrum of burst onset times (Figure 4C and 4D) - was significantly different than empirical measurements (Figure 4E and 4F). The resulting map of the parameter space of the polychronous network revealed two regions which could be distinguished by the experimentally observed distribution of burst onset times (Figure 4G). Where the delay distribution is narrow and/or the mean delay very small, sequences are based on synchronized groups of neurons, similar to a synfire chain, and thus incompatible with HVC dynamics. Above a minimum mean and SD of the delay distribution, polychronous network sequences are continuous, matching song-related activity (e.g., Figure 2E). These activity patterns are a consequence of the requirement of convergence at the postsynaptic neuron and the range of conduction delays from presynaptic neurons. In cases where there are heterogeneous delays (e.g., Figure 3C), inputs can converge onto the same postsynaptic neuron from presynaptic neurons active at different times, enabling continuous sequences. In contrast, if the range of delays is very narrow (e.g., Figure 3F), convergent inputs necessarily originate from presynaptic neurons active within this very narrow range, resulting in a sequence consisting of synchronous groups of neurons.

**Figure 4.**
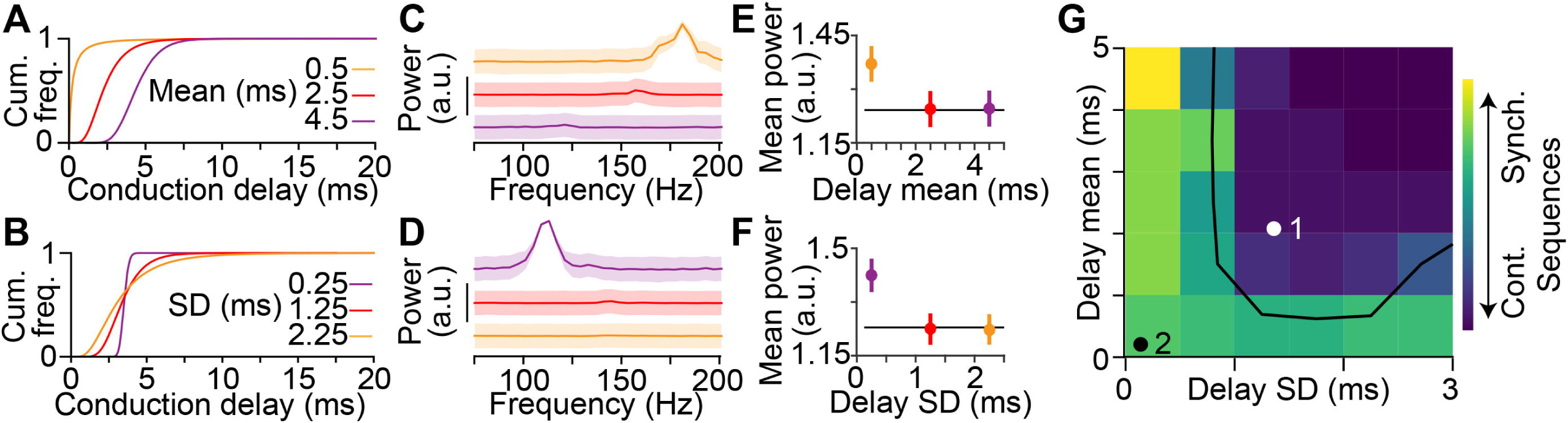
Polychronous network models support two distinct network sequence patterns depending on the parameters of the underlying delay distribution. (A) Three distributions of conduction delays with different means and identical variances (SD: 1.25 ms). (B) Three distributions of conduction delays with identical means (mean: 3.5 ms) and different variances. (C, D) Generating a polychronous network model for each distribution (C: varying the mean; D: varying the SD) allows investigating which conduction delay distributions result in continuous network sequences or sequences with synchronous groups of neurons. (E, F) Mean power in the frequency band from 75-200 Hz for the three models in (C) and (D) and the observed burst times (black line) allows measuring the degree of synchrony in the different networks. Error bars: 5^th^ and 95^th^ percentiles (bootstrap). (G) Two-dimensional parameter grid of polychronous networks with different mean and SD of delay distributions investigated here. Each grid point is colored according to the mean power of the burst onset times to determine whether the underlying network produces continuous network sequences (i.e., low power, indicated in dark blue) or sequences with synchronous groups of neurons (i.e., high power, indicated in green and yellow). White/black: Location of the models in Figure 3B (1) and 3F (2) on the parameter grid. Black line: Models to the left and below this line display sequences with synchronously active groups of neurons that are not observed in HVC (p < 0.05, bootstrap).

### Slow conduction velocity of local HVC axon collaterals

Our polychronous network model puts a strong lower bound on the conduction delays that must exist in the HVC circuit, and our value for axonal delays, based upon published mammalian conduction velocities, is close to the boundary between continuous and discrete network sequences (Figure 4G), resulting in mild, transient synchronous activity in the beginning of the sequence (Figure 3C). We therefore decided to obtain a more precise estimate of these delays through experimental observation. A direct measure of conduction delays is complicated by the fact that local unmyelinated axons are typically thin and difficult to record (Shu et al., 2006). However, HVC neurons often exhibit long-range unmyelinated axons that target the downstream song production structure (i.e., the robust nucleus of the arcopallium, RA) (Figure 1B and 5A and S2). We reasoned that we could measure the conduction velocities of these fibers and then relate long-range delays to those of the local axons within HVC. We measured the conduction velocity of action potentials in the HVC→RA projection axons by quantifying the time required for an antidromic spike initiated in RA to travel to the soma (Figure 5A, S2A and S2B) (Hahnloser et al., 2006), a path distance of 2.8 ± 0.2 mm (n = 4 reconstructions, Figure 5B). We then compared the morphological properties of descending axons – restricting our view to only the unmyelinated fibers (Figure S2C and S2D) – with another EM data set in which local axons of HVC premotor neurons were labeled (Figure 5C and 5D). Local axons were invariably unmyelinated and significantly thinner (167 ± 73 nm) than unmyelinated descending axons (446 ± 135 nm) (Figure 5D, S2E and S2F). Assuming that biophysical properties of the unmyelinated local and long-range axons are similar, we used cable properties to convert the conduction velocity measurements of the descending axons to those of local HVC collaterals (Hodgkin and Huxley, 1952; Rushton, 1951). The estimated conduction velocity of HVC axon collaterals (0.187 ± 0.035 mm/ms) enabled us to infer the propagation time for spikes to travel from the soma to different parts of the axon (Figure 5E). Using this estimate, we found delays ranging from 1 to 7.5 ms (5^th^ and 95^th^ percentiles) and up to 22 ms. This more precise estimate of the conduction velocity is considerably slower than our previous assumption and places HVC comfortably within the parameter region of polychronous networks producing continuous sequences (Figures 5F and S3).

**Figure 5.**
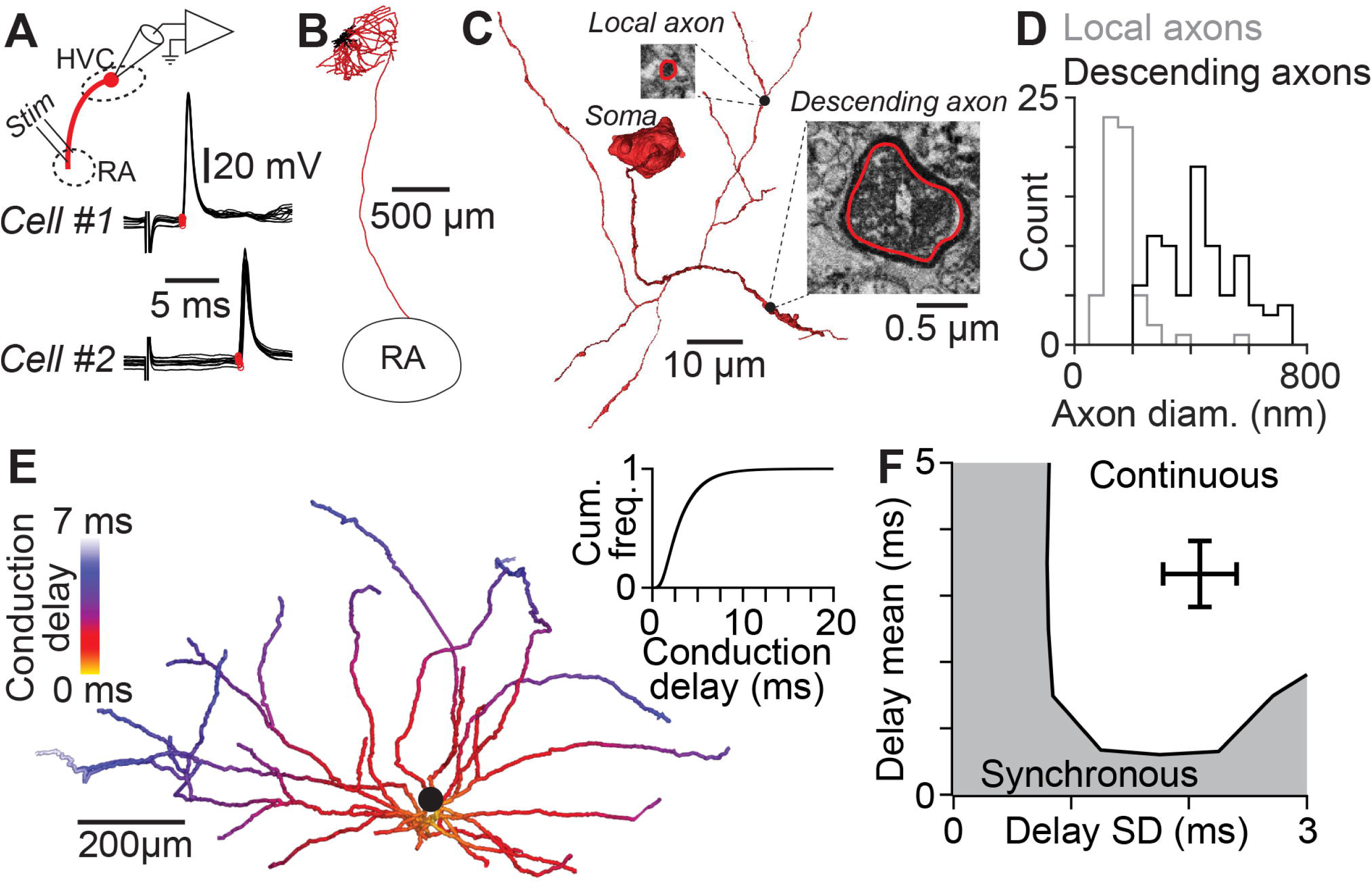
Local HVC premotor neuron axons have conduction delays supporting continuous sequences. (A) Antidromic stimulation of HVC premotor neuron axons and whole-cell recording at the soma allows precise measurement of conduction times along the descending axon. Stimulus artifact is blanked for visualization. (B) Example reconstructions of the projection axon of a premotor neuron connecting HVC and RA. (C) 3D reconstruction of the soma and proximal axons of a retrogradely labeled HVC premotor neuron from an SBEM image stack. Insets: EM micrographs of labeled axons. (D) Unmyelinated axons in the HVC-RA fiber tract have larger diameters than local (unmyelinated) collaterals of HVC premotor neurons. (E) Measurement of the conduction velocity along unmyelinated long-range axons allows a precise estimate of the conduction delays along thin unmyelinated local axon collaterals of HVC premotor neurons based on 22 reconstructions of local axon collaterals. (F) The distribution of conduction delays along HVC premotor neuron axons supports a polychronous network model generating continuous sequences (see Figure 4G).

### Polychronous network organization explains HVC spatiotemporal activity

To this point, we have demonstrated that the polychronous model given our measured experimental constraints can explain the temporal structure of HVC function at a network (i.e., continuous representation) as well as at a cellular (i.e., axonal conduction delays) level. We next asked whether this underlying circuit structure can predict other aspects of song-related HVC function. To accomplish this, we returned to our local axonal collateral reconstructions, and we placed synapses at specific locations within the axonal field that fit a variety of conduction delay distributions (Figure 6). For instance, in cases in which the conduction delays were long but exhibited a low variance, synapses were clustered on distal axons (Figure 6A). We can also look at cases in which the means of the conduction delays were low, across two variance conditions (Figures 6B and 6C). We compare these possibilities against a scenario that matches our experimental observations in which the mean and variance were both relatively high (Figure 6D). Because of the differential placement of synapses within the axonal field, each model should result in a different prediction concerning the spatiotemporal pattern of activity in HVC during singing (Graber et al., 2013; Markowitz et al., 2015; Peh et al., 2015). We performed 2-photon imaging of GCaMP6-expressing projection neurons during singing to measure the activity of 182 putative premotor neurons (see Methods), combining new observations with a previously published data set (Katlowitz et al., 2018; Picardo et al., 2016) (Figure 6E). Using an established algorithm, we precisely estimated burst onset times with a temporal resolution of ~10 ms (Picardo et al., 2016) and related these values to the relative spatial position for each neuron (Figure 6F). We excluded pairs in which the difference in burst times was greater than 20 ms and therefore unlikely to be driven by monosynaptic connections. In the remaining cases, sequentially active neuron pairs were found over a wide range of relative locations (178 ± 102 µm, mean ± SD), from immediately adjacent (~10 µm) to much longer distances (~500 µm, or approximately one third of the maximum extent of HVC) (Figure 6G and 6H). We then compared the spatial location of sequentially active pairs against the predictions of our previously stated models. Whereas three models predicted a high degree of spatial clustering (Figure 6A–6C), the model based on our empirically measured delay distributions was more spatially dispersed (Figure 6D), matching the functional data (Figure 6I). We conclude that a polychronous network sequence based on conduction delays observed in HVC results in a spatial organization that closely resembles observations of HVC network activity during song.

**Figure 6.**
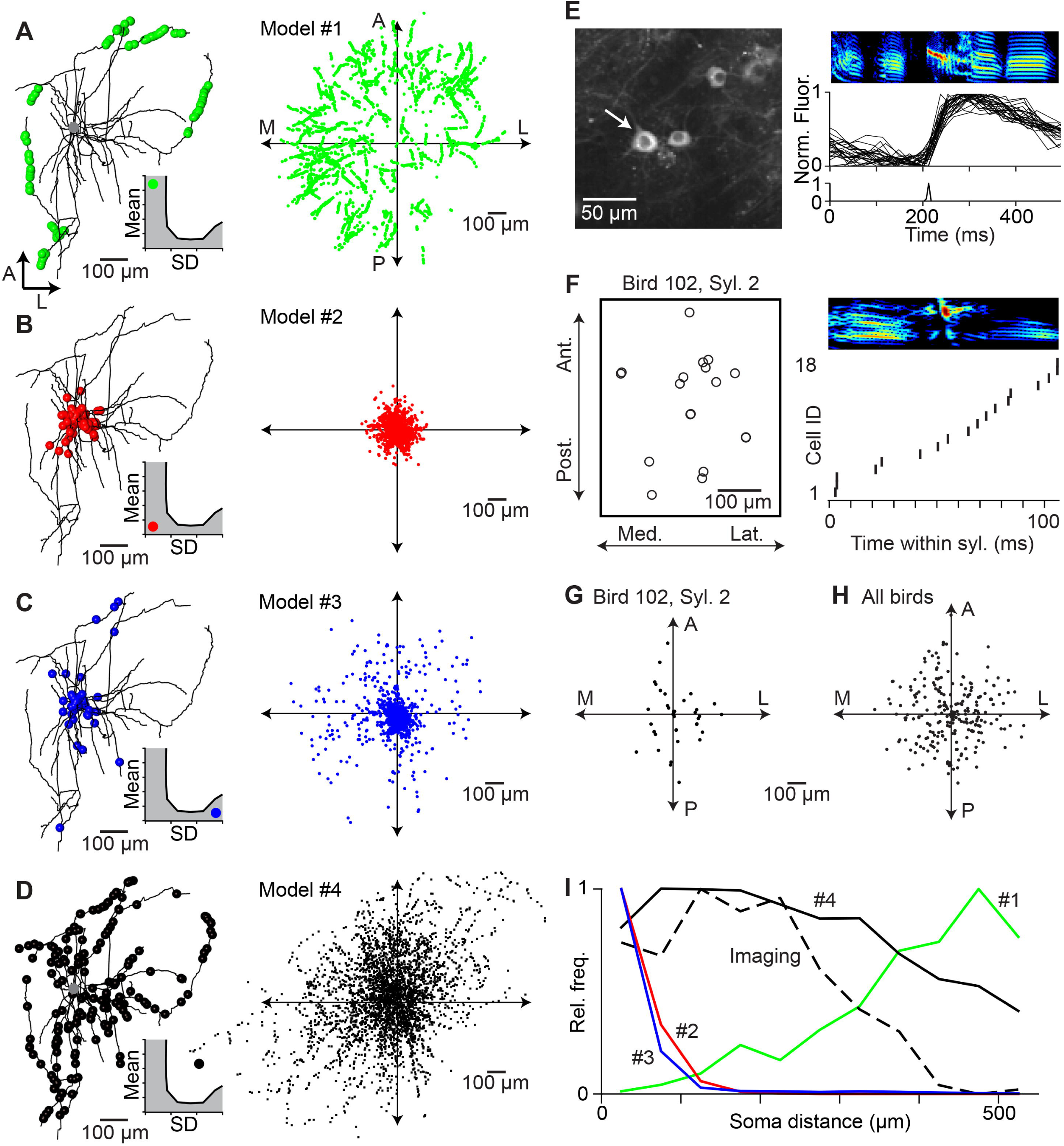
Spatial organization of a polychronous network matches HVC projection neuron activity during singing. (A-D) Left: Distribution of active synapses along the local axonal collaterals of a premotor neuron for a given delay distribution (inset; A: mean / SD: 4.5 / 0.25 ms; B: 0.5 / 0.25 ms; C: 0.5 / 2.75 ms; D: 3.3 / 2.1 ms). Right: Distribution of active synapses relative to the soma based on the local axonal collaterals of 22 HVC premotor neurons. (E) 2-photon calcium imaging of song-related bursting activity in HVC. Left: Example image of GCaMP6s-labeled somata. Right: Spectrogram of song motif (top), aligned normalized fluorescence traces of the neuron highlighted in left panel (center), and estimated burst onset time (bottom). (F) Left: Soma locations of 18 neurons active within the same syllable, projected onto the horizontal plane. Right: Estimated burst onset times of the same neurons within the syllable. (G) Relative soma locations of putatively connected neurons in F (i.e., burst onset times within 20 ms of each other). (H) Relative soma locations of putatively connected neurons in nine birds. (I) Radial distribution of putative postsynaptic neurons in (H) (dashed line) and radial distribution of active synapses predicted by the four network models in A-D (solid lines).

### Delay distributions are conserved from songbird to mammalian neocortex

We have demonstrated an important impact of local axonal delays on the timing and structure of network activity within HVC of the zebra finch. Given the extraordinarily slow axonal conduction velocity in HVC compared with known values measured in a variety of different circuits (Figure 7A), it remains unclear whether such delays will play a role within those networks or whether this solution is simply a specialization within zebra finch HVC. To begin to examine this issue, we analyzed the local collaterals of 14 spiny neurons in Layer 4 of rat somatosensory cortex (Figure 7B) (Narayanan et al., 2015). When we measured the pathlength from the soma to different locations along the axon, we found that the entire size of the axonal field was considerably larger than that of HVC premotor neurons (Figure 7C). Surprisingly, when we estimated conduction delays – accounting for both the discrepancies in conduction velocity and pathlength – we find that the range of these values is identical in both cell classes (L4: 3.4 ± 2.3 ms, mean ± SD; HVC: 3.3 ± 2.1 ms, Figure 7D). Therefore, significant conduction delays exist within the rodent neocortex, potentially playing an important computational role within that circuit.

**Figure 7.**
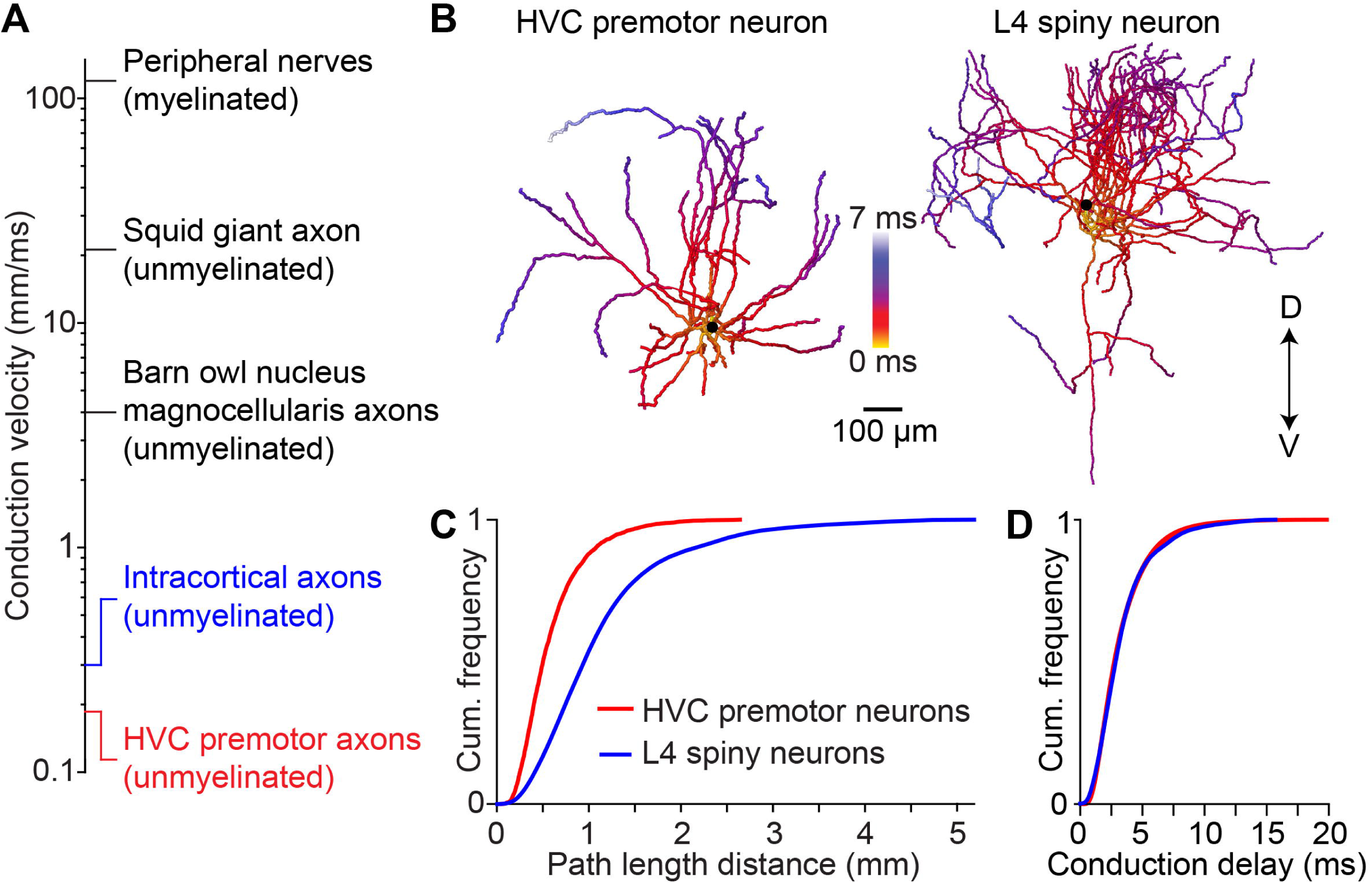
Conduction delays of neocortical axonal arbors. (A) Axonal conduction velocity measurements from the peripheral to the central nervous system. Peripheral nerves: (Hursh, 1939); squid giant axon: (Hodgkin and Huxley, 1952); nucleus magnocellularis axons: (Carr and Konishi, 1990); intracortical axons: (Hirsch and Gilbert, 1991; Shu et al., 2007). (B) Example reconstructions of an HVC premotor neuron axon and a L4 spiny neuron axon from rat somatosensory cortex. Conduction delays estimated based on conduction velocity measurements in (A). D/V: dorsal/ventral. (C, D) Distributions of axonal pathlengths (C) and resulting conduction delays (D) for 22 HVC premotor neuron axons and 14 L4 spiny neuron axons (Narayanan et al., 2015).

## DISCUSSION

Using a range of modeling and experimental approaches, we investigated how local excitatory circuits can give rise to convergent synaptic input underlying sequential activity in the zebra finch song system. We provided three independent lines of evidence supporting a central role for slow and heterogeneous axonal conduction delays in HVC sequence generation. First, network modeling revealed that delays are required to generate the continuously active population sequences observed in HVC. Second, the delays predicted by the network model match empirically measured values for local HVC axon collaterals. Third, the spatiotemporal patterns of HVC activity observed during singing matches the predictions from our model. As a result, we propose that the core circuit for sequence generation in HVC consists of an asynchronous feedforward network based on a variety of conduction delays to generate a continuous neural sequence. Notably, previous theoretical work has demonstrated that axonal delays in synfire chain networks can enable more continuously active network sequences by ‘skipping’ connections between different groups, therefore enabling so-called ‘synfire braid’ networks (Bienenstock, 1996) which function similarly to polychronous networks. Future studies will elucidate how these sequences are started (Figure S3) (Andalman et al., 2011; Danish et al., 2017; Galvis et al., 2018) as well as the role of other circuit elements, such as local circuit interneurons, in this process (Gibb et al., 2009; Jin et al., 2007; Kosche et al., 2015; Yildiz and Kiebel, 2011).

In this study, we provide the first demonstration of a biological neural network implementing the polychronous circuit structure. We find that this network is capable of producing sustained sequential activity over behaviorally relevant timescales (Itskov et al., 2011), exceeding the predictions from the original model. The continuous neural sequences arising from this model can represent any moment in time, thereby greatly increasing the resolution of the HVC premotor clock and facilitating the placement of descending motor commands at any time point during the song. While the present model assumes that synaptic connections are made within a synchronous time window, another feature of polychronization is the potential to self-organize such connectivity patterns through spike timing-dependent plasticity mechanisms (Gerstner et al., 1996; Izhikevich, 2006), and future work will determine the relevance of axonal delays for assembly of HVC circuits during song learning (Fiete et al., 2010; Jun and Jin, 2007; Okubo et al., 2015).

We found that slow axons within a local circuit are critical for continuous sequence generation. We estimate that conduction delays are significantly larger than those afforded by other biophysical parameters, such as synaptic delays (Sabatini and Regehr, 1996) (~0.1 ms) and postsynaptic integration time (Long et al., 2010) (~5 ms during singing). Notably, the dendrites of HVC premotor neurons are highly compact (i.e., a spatial extent of less than 200µm) (Benezra et al., 2018) and are unlikely to contribute significantly, although to date no direct measurements of their electric signaling properties exist (Magee and Cook, 2000; Williams and Stuart, 2002). Therefore, approximately half of the total elapsed time of the HVC sequence could be attributed to local axonal conduction, a comparatively inflexible process that may underlie the behavioral stereotypy inherent in the adult zebra finch song (Lombardino and Nottebohm, 2000) by rendering the circuit less sensitive to perturbations (Hamaguchi et al., 2016; Swadlow et al., 1981). We anticipate that further studies will clarify the impact of additional time delays introduced along the pathway from HVC to syringeal motorneurons (Figure S2A) in converting the HVC code to behavior and the extent to which the axonal properties themselves (e.g., myelination status or axial diameter) can be modified through experience. Furthermore, although our model does take into account a range of axonal diameters for local collaterals, the detailed effects of precise axonal morphology – e.g., possible failures at branch points (Swadlow et al., 1980) – remain to be explored.

The idea that axons contribute to information processing in neural circuits has long been explored for long-range connections between different brain areas (Carr and Konishi, 1988; Innocenti et al., 1994; Salami et al., 2003; Sugihara et al., 1993). For instance, in the brainstem of the barn owl, axons carrying sound information from both ears form precisely tuned and spatially organized ‘delay lines’ (Jeffress, 1948) necessary for detecting minute interaural time differences (Carr and Konishi, 1988). In contrast, the role of axonal delays within local microcircuits is often disregarded (Budd et al., 2010), possibly because of the technical challenges involved in obtaining reliable estimates of conduction velocity in local circuits. In this study, we find that the premotor song production structure HVC in the zebra finch uses ‘delay lines’ within a local circuit to generate reliable sequences of activity during song (Katlowitz et al., 2018). We do not yet know whether this specialization is unique to circuits in which a high degree of temporal precision is required or more broadly found in other networks, including those capable of more flexibility. Although we find that conduction delays along intracortical axons in a rodent neocortical area are likely to be comparable to those reported here in zebra finch HVC, the extent to which these collaterals may support persistent activity that has been observed within this region so far remains unexplored (Sachidhanandam et al., 2013). Further work in other circuits can establish whether this delay distribution represents a universal scaling law (Buzsaki and Mizuseki, 2014; Liewald et al., 2014; Miller, 1996) across different species, brain regions, cell types, etc., or whether these local delays are specially tuned for the requirements of each unique case. Overall, our results suggest that in addition to defining the static architecture of neural networks (Denk et al., 2012; Plaza et al., 2014; Seung, 2012), functional properties of axons within local circuits can also control the space of neural activity patterns.

## ACKNOWLEDGEMENTS

We thank Dmitriy Aronov, György Buzsáki, Dmitri Chklovskii, Yarden Cohen, Annegret Falkner, Dan Levenstein, Cengiz Pehlevan, Alex Reyes, John Rinzel, Richard Tsien, and members of the Long laboratory for comments on earlier versions of this manuscript. We thank the Fee laboratory for providing extracellular recordings of HVC projection neurons and Marcel Oberlaender for the reconstructions of L4 neurons. We thank NYU Langone’s Microscopy Laboratory for assistance with electron microscopy. We also acknowledge helpful conversations with Asohan Amarasingham, Yoram Burak, Dina Obeid, and Kanaka Rajan.

## AUTHOR CONTRIBUTIONS

R.E. and M.A.L conceived the study and designed the experiments; S.E.B., M.A.L., M.A.P, F.M. and J.K. conducted the research; R.E., K.A.K., S.E.B., M.A.P., F.M., J.K., and M.A.L. performed data analyses; Y.T. and D.Z.J. developed the theoretical model; R.E., Y.T., K.A.K., and M.A.L. created the figures; R.E. wrote the initial draft of the manuscript; R.E., D.Z.J., and M.A.L. edited and reviewed the final manuscript. R.E., M.A.L., and D.Z.J. acquired funding; M.A.L. and D.Z.J. supervised the project.

## STAR METHODS

### CONTACT FOR REAGENT AND RESOURCE SHARING

Further information and requests for resources and reagents should be directed to and will be fulfilled by the Lead Contact, Michael Long.

### EXPERIMENTAL MODEL AND SUBJECT DETAILS

We used adult (>90 days post hatch) male zebra finches (*Taeniopygia guttata*) that were obtained from an outside breeder and maintained in a temperature- and humidity-controlled environment with a 12/12 hr light/dark schedule. All animal maintenance and experimental procedures were performed according to the guidelines established by the Institutional Animal Care and Use Committee at the New York University Langone Medical Center.

## METHOD DETAILS

### Surgeries

Surgical procedures for retrograde labeling of HVC premotor neurons, viral injections, chronic cranial window implantation for 2-photon imaging, and *in vivo* whole cell recordings, have previously been described in detail (Kornfeld et al., 2017; Long et al., 2010; Picardo et al., 2016). Briefly, animals were anesthetized (1-3% isoflurane in oxygen) and the scalp was cut to expose the entire skull. To label HVC premotor neurons for electron-microscopic imaging, a biotinylated dextran (BDA-dextran, MW: 3,000; Invitrogen) was injected into RA. Birds were allowed to recover for three days to allow retrograde labeling. For virus injections, a craniotomy was made over HVC and either AAV9.Syn.GCaMP6s.WPRE.SV40 (Penn Vector Core) or a 1:1 mix of AAV9.CamKII0.4.Cre.SV40 and either AAV9.CAG.Flex.GCaMP6f.WPRE.SV40 or AAV9.CAG.Flex.GCaMP6s.WPRE.SV40 was injected using an oil-based pressure injection system (Nanoject 3, Drummond Scientific). After injections, a cranial window was implanted over the craniotomy. For antidromic activation of HVC premotor neurons, a bipolar stimulation electrode was implanted into RA. Then, a craniotomy was made over HVC for whole-cell recordings during sleep.

### *In vivo* whole-cell recordings

Whole-cell recordings of HVC premotor neurons during sleep were made with glass electrodes (~5-8 MΩ) using previously described techniques(Long et al., 2010). Briefly, pipettes were advanced with positive pressure while monitoring resistance. Proximity to neurons was indicated by a resistance increase, and pressure was released to allow formation of a gigaseal. The membrane was ruptured by applying negative pressure. After establishing whole-cell configuration, series resistance was compensated and the membrane potential recorded in current-clamp mode. Recordings with series resistance vales greater than 30 MΩ and resting membrane potentials more depolarized than −60 mV were discarded. HVC premotor neurons were identified by evoking antidromic action potentials upon stimulation of RA (10-100 µA amplitude, bipolar stimulation of 0.2ms duration).

### 2-photon calcium imaging

The procedures for 2-photon calcium imaging of HVC neurons during singing have been described previously (Katlowitz et al., 2018; Picardo et al., 2016). Briefly, birds were first trained to perform directed singing in the head-fixed configuration upon presentation of a female bird using operant conditioning with a water reward (Picardo et al., 2016). Once the behavior was learned, virus injections and cranial window implantation were performed. 2-photon calcium imaging was carried out using a resonant scanning system (Thorlabs) at a frame rate of 28.8 Hz and a 16x water-immersion objective (NA 0.8, WD 3 mm; Nikon). We acquired singing behavior using an omnidirectional microphone (Audio*-*Technica) digitized at 40 kHz (Digidata 1550, Molecular Devices). Motif-related image data were temporally aligned offline by linearly warping to manually annotated reference points within the song. Fluorescence traces were extracted from manually drawn ROIs (ImageJ) on temporally aligned and motion-corrected (Miri et al., 2011) image stacks. Frame times were defined for each neuron as the mean time point that the laser reached the ROI as a function of vertical scanning location. Last, burst onset times were deconvolved from the raw traces using a Markov Chain Monte Carlo inference approach with an average uncertainty of ~10 ms (Picardo et al., 2016; Pnevmatikakis et al., 2016). These analyses were performed using custom Matlab scripts (Mathworks).

### Extracellular recordings during singing

We reanalyzed a previously reported data set of burst times from HVC neurons during singing (Lynch et al., 2016; Okubo et al., 2015). The data set contained extracellular recordings obtained in two adult birds (i.e., ≥103 d.p.h.) and three young adult birds (i.e., ≥59 d.p.h.) with a stable motif. Single units were identified as HVC projection neurons (i.e., projecting to nucleus RA or along the anterior forebrain pathway to Area X) by antidromic stimulation or as putative HVC projection neurons based on low spontaneous firing rate (i.e., <1 Hz) and sparse bursting activity during singing. Individual song motifs and the accompanying neural activity were time-warped to syllable on- and offsets. Then, the firing rate was computed in 1 ms bins and smoothed with a 9 ms wide sliding window. A ‘burst window’ was defined as a period where the smoothed firing rate exceeded a threshold of 10 Hz. To define the burst onset more precisely, each spike falling into the ‘burst window’ was replaced with a 5 ms wide ‘spike interval’ starting at the spike time. The burst onset time is defined as the earliest time point in a ‘burst window’ where at least three ‘spike intervals’ from different song renditions overlap.

### Histological procedures

For serial block-face electron-microscopic (SBEM) imaging, perfusion and histology was performed as described in detail previously (Kornfeld et al., 2017). For transmission electron-microscopic imaging, the protocol used for SBEM imaging was slightly modified as follows. After the bird was transcardially perfused, the brain was removed from the skull and post-fixed overnight (Kornfeld et al., 2017). The brain was then cut into 100 µm thick slices using a vibratome (Leica VT1000S). Residual peroxidase activity was suppressed by soaking the sample in 3% H_2_O_2_ for 20 min before labeling the sample with an avidin-peroxidase complex and DAB. A slice containing clearly visible stained fibers from HVC to RA was carefully unmounted by immersing the microscope slide into PB. After washing with PB, the samples were post fixed in 1% OsO_4_ for 2 hours, block stained with 1% uranyl acetate for 1 hour, dehydrated in ethanol and embedded in EMbed 812 (Electron Microscopy Sciences, Hatfield, PA). Semi-thin sections were cut at 1 µm and stained with 1% toluidine blue to find the previously identified area of interest containing fibers from HVC to RA. In each sample, 20 serial ultrathin sections with 100 nm thickness were cut, mounted on slot copper grids, and stained with uranyl acetate and lead citrate.

### Transmission-electron microscope imaging

Stained grids were examined under a Philips CM-12 electron microscope (FEI; Eindhoven, The Netherlands) and photographed with a Gatan (4k x 2.7k) digital camera (Gatan, Inc., Pleasanton, CA). Samples were imaged at a series of increasing magnifications (i.e., ranging from 3,400x to 66,000x magnification) to allow identification of fiber tracts and ultimately individual fibers within these tracts. Diameter measurements of unmyelinated projection axons were made on images with a magnification of at least 40,000x.

### Axon diameter measurements

All light micrographs used for illustration of local and descending axons were captured using a Zeiss AxioObserver Inverted. We acquired images of descending HVC premotor neuron axons from ultrathin sections using a transmission electron microscope (see above). Unmyelinated descending axons were identified based on dark DAB labeling in EM micrographs. Myelinated axons were identified morphologically by presence of multiple, closely wrapped membrane layers (i.e., myelin sheaths). Diameters were measured along the shortest axis of the circumference of each axon (i.e., if the axon was cylindrical, this corresponds to the diameter of the cylinder irrespective of sectioning angle) (Figure S2F). Diameters of local HVC premotor neuron collaterals were measured using a previously reported data set acquired using serial block-face EM (Kornfeld et al., 2017) with a voxel size of 11 x 11 x 29 nm^3^ containing HVC premotor neurons labeled by injection of a tracer (BDA-dextran) into RA. Diameters of randomly selected locations along labeled local axon collaterals were measured by determining the image plane that was closest to the orthogonal plane defined by the axon and measuring the axon diameter in that plane.

### Estimating synapse locations along axons

We estimated the possible locations of synapses between HVC premotor neurons along local axons by combining results from previously reported anatomical data sets (Benezra et al., 2018; Kornfeld et al., 2017). We used a database of 22 reconstructions of local axon collaterals of *in vivo* labeled HVC premotor neurons (Benezra et al., 2018) to determine possible synapse locations irrespective of the postsynaptic target along the local axons of each neuron by sampling points along the reconstructed axons at the average distance between synapses, which has previously been determined using EM measurements along HVC premotor neuron axons (Kornfeld et al., 2017). We then estimated which of these possible synapse locations could target other HVC premotor neurons. To do so, we fitted previous EM measurements of the relative frequency of HVC premotor neurons as the postsynaptic target as a function of distance to the presynaptic soma with a sigmoidal function. For each estimated synapse location along the reconstructed axons, we then computed the synapse-soma distance and placed a synapse at this location with a probability equal to the corresponding relative frequency. Finally, we computed the pathlength distance between each of these possible premotor synaptic locations and the soma (i.e., the shortest path along the axon connecting these two points).

Synapse locations along HVC premotor neuron axons for a given delay mean and SD were estimated as follows. Points along the reconstructed axon were grouped according to their pathlength distance to the soma into 50 µm bins. If successive points in the reconstruction had an interval of more than 1 µm, additional points were inserted at 0.5 µm intervals using linear interpolation (i.e., leaving the pathlength unchanged). Next, the log-normal delay distribution with given mean and SD was converted to a pathlength distribution by multiplication with the axonal conduction velocity of local HVC premotor neuron axons (i.e., 0.187 mm/ms). For each neuron, we generated N_Syn_ * L_Neuron_ / L_Avg_ samples from this log-normal distribution. Here, N_Syn_ = 170 is the average number of synapses made by each HVC premotor neuron onto other premotor neurons, L_Neuron_ is the total axonal pathlength of this specific premotor neuron and L_Avg_ is the average axonal pathlength across all 22 premotor neurons (Benezra et al., 2018). Samples beyond the maximum pathlength distance to the soma were repeated. A histogram of these samples with a bin width of 50 µm was computed. For each 50 µm bin, points along the reconstruction in the corresponding pathlength bin were randomly sampled until the number of elements in this bin of the histogram was reached and a synapse was placed at the location of each sampled points along the reconstruction.

### Estimating axonal conduction delays

The first estimate of conduction delays along local HVC premotor neuron axons was obtained by measuring the pathlength from the soma to points along the axon spaced at the mean inter-bouton interval of HVC premotor neuron axons (10.5 μm) (Kornfeld et al., 2017). Combining these measurements from 22 reconstructions of premotor neuron axons reported previously (Benezra et al., 2018) resulted in and average distribution of pathlengths. We then converted these pathlengths into a conduction delay distribution by multiplying each pathlength distance with a value of 0.3 mm/ms for the conduction velocity of unmyelinated neocortical axons. A log-normal distribution described the shape of the conduction delay distribution well (least-squares fit R^2^ = 0.9988). We therefore used mean and standard deviation of a log-normal distribution to parameterize the conduction delays for the polychronous network models.

Conduction delays along L4 spiny neuron axons were estimated in the same way. We measured the distribution of pathlength distances of 14 complete reconstructions of the intracortical axonal arbor of L4 neurons labeled *in vivo* (Narayanan et al., 2015) and multiplied pathlengths by a conduction velocity of 0.3 mm/ms to obtain the conduction delay distribution.

The conduction time along long-range axons from HVC to RA was measured from whole-cell membrane potential recordings of HVC premotor neurons as the difference between the onset time of antidromic stimulation in RA and action potential onset. The action potential onset was defined by calculating the second derivative of the membrane potential between 0-20 ms after stimulation and determining the first upward threshold crossing, where the threshold was set as the minimum of either 3 standard deviations of the second derivative or 400 mV/ms^2^. To determine the threshold between putative groups of conduction delays, we used k-means clustering with two groups. The pathlength of the long-range axon of HVC premotor neurons was measured from the soma to the first bifurcation of the axon as it entered RA. The average conduction velocity of unmyelinated descending axons was calculated by dividing the average descending pathlength by the average conduction time of the second mode of the conduction time distribution measured as described above. We then used a simple biophysical model relating the diameter of unmyelinated axons to conduction velocity (Hodgkin and Huxley, 1952; Rushton, 1951):

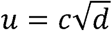

Here, *u* is the conduction velocity, *d* the axon diameter, and *c* a constant. We determined *c* using the average conduction velocity and average diameter of putative unmyelinated descending axons and assumed that this constant is the same for unmyelinated local axons of HVC premotor neurons (i.e., that the basic biophysical properties underlying action potential propagation are the same). We then calculated a distribution of conduction velocities given the observed distribution of diameters of local axonal projections. To estimate the distribution of conduction times to synapses onto other HVC premotor neurons, we used a Monte Carlo simulation approach. We stepped through all possible synapse locations along the set of reconstructed axon morphologies of HVC premotor neurons. For each possible location, we calculated the distribution of conduction times to that location given the pathlength to the soma and the estimated distribution of local conduction velocities. We then randomly selected one of the possible conduction times and randomly assigned it as a synapse onto other HVC premotor neurons based on EM measurements of premotor synapse density for each location relative to the soma (Kornfeld et al., 2017). We ran 100 Monte Carlo simulations to obtain a robust estimate of the resulting conduction time distribution to other HVC premotor neurons.

### Frequency analysis of burst onset times

For each syllable in the electrophysiology data set, we determined the syllable length and number of burst onset times occurring during the syllable. In our modeling effort (see Figure 2F, 3D and 3G), we simulated possible burst onset time distributions by sampling random numbers distributed in time according to the burst density of the model, while preserving the distribution of syllable lengths from the experimental data sets and the number of burst onset times observed during each syllable. For each syllable, we then defined the power spectral density P_s_ of the burst times as the absolute magnitude squared of the discrete Fourier transform evaluated at frequencies *f* between 1 and 300 Hz, in increments of 4 Hz:

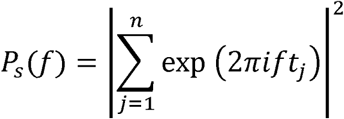

Here, *n* is the number of bursts in the syllable, and *t_j_* the burst onset time of the *j^th^* burst. We then calculated the mean power spectrum across all syllables. In order to obtain a reliable estimate of the predicted power spectrum of each model and its uncertainty, we repeated this procedure 10,000 times, and computed the mean and standard deviation at all evaluated frequencies. For comparison with experimental data, we set the confidence interval as ±3 standard deviations.

### Neuron and synapse models

HVC premotor neurons were modeled as a two-compartment model with a dendritic and somatic compartment (Long et al., 2010). Current injection at the soma triggers sodium channel-dependent action potentials, while current injection (and synaptic input) to the dendrite compartment triggers an all-or-none calcium spike, which in turn triggers a high-frequency burst of four action potentials at the soma. Ion channels were modeled using the Hodgkin-Huxley formalism. All model parameters are identical to our previous work (Long et al., 2010), except for the following differences: R_c_ = 130 MΩ, G_s,L_ = 0.05 mS/cm^2^, τ_c_ = 15 ms. Conductance-based excitatory synapses were modeled according to ‘kick-and-decay’ dynamics. Upon synaptic release, the synaptic conductance was increased by G_syn_, followed by an exponential decay with time constant τ_syn_ = 5 ms. The weight G_syn_ of individual synapses was drawn from a uniform distribution [0, G_max_], with G_max_ set to 0.05 mS/cm^2^, a value that leads to a unitary EPSP of ~4 mV at the soma (i.e., the average EPSP amplitude is 2 mV (Mooney and Prather, 2005)).

### Synfire chain network assembly

The synfire chain network model of HVC premotor neurons was constructed by sequentially connecting 117 nodes with 170 neurons in each node, corresponding to 19,890 neurons in the entire network. Neurons in each node of the synfire chain were connected in a feed-forward all-to-all manner to the neurons in the subsequent node.

### Feedforward polychronous network assembly

The feedforward polychronous network (Izhikevich, 2006) was assembled in an iterative process. The algorithm was designed to enforce synchrony of the synaptic inputs to the postsynaptic neurons. The timing of presynaptic bursts must be such that the spikes arrive at the postsynaptic neuron within a narrow time window (discussed below), taking into account the different axonal conduction delays. The number of connections that each neuron can receive was limited to 170 (Kornfeld et al., 2017). The axonal delays of these connections were based on observed delay distributions.

Each iteration consisted of three steps (Figure S1A): *(i)* simulation of network dynamics to determine the burst onset times of all neurons in the network at the current iteration; *(ii)* adding feedforward connections constrained by a given conduction delay distribution between ‘source neurons’ (i.e., presynaptic neurons) and ‘target neurons’ (i.e., potential postsynaptic targets which do not form outgoing connections in the current iteration); *(iii)* adding additional neurons into the network. As a result, the feedforward network grows in size and the corresponding population sequence in duration during this iterative process (Figure S1B). In *<u>step (i)</u>* of each iteration, network dynamics were simulated by activating a set of 200 predefined ‘starter neurons’ and recording burst onset times of all active neurons in the network. In *<u>step (ii)</u>*, new feedforward synaptic connections between ‘source’ and ‘target neurons’ were added. First, N_new_ neurons were moved from the set of ‘target neurons’ in the previous iteration to the set of ‘source neurons’. Specifically, these were the ‘target neurons’ whose simulated burst onset times were within a 2 ms window from the earliest simulated burst onset time of all ‘target neurons’ (Figure S1C). We then generated a ‘synaptic pool’ (i.e., a set of conduction delays τ_ax_) of size N_out_ * N_new_ (Nout = 170)(Kornfeld et al., 2017) by sampling from the given distribution of conduction delays. We iterated over all ‘target neurons’ ordered according to the number of synaptic inputs, starting with the smallest number. For each ‘target neuron’, we randomly selected a ‘source neuron’ that fulfilled the polychronization requirement | t_target_ – τ_int_ – τ_ax_ – t_source_ | ≤ τ_sync_ using a suitable conduction delay τ_ax_ from the ‘synaptic pool’ (Figure S1D; i.e., requiring that all synaptic inputs to the ‘target neuron’ arrive within a synchronous time window 2*τ_sync_ (here: time window of 1 ms). t_source_ is the burst onset time of the ‘source neuron’, τ_int_ is the average integration time constant of HVC premotor neurons from onset of the synaptic inputs to burst threshold (set to 5 ms) (Long et al., 2010), and t_target_ is the burst onset time of the ‘target neuron’. If there were multiple τ_ax_ allowing a connection between the ‘source neuron’ and the ‘target neuron’, the one minimizing the quantity | t_target_ – τ_int_ – τ_ax_ – t_source_ | was selected (i.e., only one synapse was placed between a pair of ‘source’ and ‘target neurons’). After placement, this synaptic connection was removed from the ‘synaptic pool’. If the number of synaptic inputs to the ‘target neuron’ reached 170 or no connection from the ‘source neurons’ could be made given the conduction delays in the ‘synaptic pool’, it was not considered as a ‘target neuron’ anymore. In *<u>step (iii)</u>*, neurons were added to the network in order to increase the network from the set of starter neurons to its final size. This step was taken in case there were no more ‘target neurons’ before the ‘synaptic pool’ was exhausted. In this case, the set of ‘target neurons’ was restored to its state at the beginning of the iteration. A new ‘target neuron’ (i.e., without any existing incoming or outgoing synaptic connections) was added to the network by placing a synaptic connection with a randomly selected conduction delay τ_ax_ from the ‘synaptic pool’ originating from one of the N_new_ ‘source neurons’ added to the network in this iteration. The putative burst onset time of the new ‘target neuron’ was defined as: t_new_ = t_source_ + τ_ax_ + τ_int_ (Figure S1E). All other synaptic connections placed in this iteration were removed from the network; the associated conduction delays were moved back into the ‘synaptic pool’; and *steps (ii)* and *(iii)* were repeated until the ‘synaptic pool’ was empty. Then, the next iteration was started, and this process repeated until all 20,000 HVC-RA neurons were incorporated into the network. To investigate the effect of conduction delays on sequence generation, we used different conduction delay distributions during network assembly. The delay distribution in the completely assembled network matched the distributions based on observed delays (Figure S1F).

### Simulations

During simulations, HVC premotor neurons received additional independent white noise input currents to their somatic and dendritic compartments with zero mean and amplitudes A_soma_ = 0.1nA and A_dendrite_ = 0.2nA, leading to fluctuations of the somatic membrane potential with a standard deviation of 4.2 mV (Long et al., 2010). To account for the white noise currents, the HVC premotor neuron models were treated as a system of stochastic differential equations and solved using the AN3D1 weak 3^rd^ order method (Debrabant, 2010). The simulation time step was set to 0.02 ms.

Each simulation was started by activating the set of 200 ‘starter neurons’ using an excitatory conductance kick with amplitude 300 nS exponential decay with time constant 5 ms (i.e., simulating synchronous synaptic input). This input was delivered to the ‘starter neurons’ either synchronously, uniformly distributed over a 7 ms window, or randomly within a 10 ms window. In order to minimize transient effects of this activation procedure, the first 50 ms of simulated activity were discarded. Network activity patterns after this transient period were qualitatively similar between the different activation procedures. To generate burst densities, we ran 50 simulations, recorded the burst onset time of each neuron (i.e., the time where the membrane potential at the soma crosses 0 mV for the first time during a burst) and calculated the average number of bursts in 0.75 ms bins.

## QUANTIFICATION AND STATISTICAL ANALYSIS

All statistical details of experiments can be found in figure legends and the Results section, including the statistical tests used, exact value of n, what n represents (e.g., number of animals, number of cells, etc.), definition of center, and dispersion and precision measures (e.g., mean, median, SD, SEM, confidence intervals). Significance was defined at a level of 0.05. Normal distribution of data was not assumed. No data were excluded from analysis. Statistical calculations were performed using MATLAB R2016a.

## DATA AND SOFTWARE AVAILABILITY

Software and documentation required for setting up and running simulations of the synfire chain and polychronous network models can be downloaded from: https://psu.box.com/s/55gh5tjgpvcxikel4wjfkzxdwyc0s7x4

## SUPPLEMENTAL FIGURE LEGENDS

**Figure S1. Polychronous network assembly. Related to Figure 3 and Figure 4.** (A) Algorithm for polychronous feedforward network assembly. (B) During each iteration, neurons are added to the network such that they are active at the end of the current network sequence. (C) In each iteration, the sets of source and target neurons are updated according to simulated burst onset times in the current state of the network. (D) Illustration of synapse placement based on polychronous principle. (E) Illustration of placement of synapses onto neurons newly added into the network. (F). Top: Observed delay distribution and log-normal fit. Center, bottom: Mean and SD of the delay distributions in assembled networks match the parameters of the input log-normal distributions. Dashed: identity line.

**Figure S2. Estimation of conduction velocity along unmyelinated axons. Related to Figure 5.** (A) Conduction delay measurements along the HVC➔RA projection axon of 40 projection neurons. Group membership was defined based on k-mean clustering with two groups. (B) Identification of ‘dashed’ (left; group *i*) or ‘continuous’ (right; group *ii*) labeling of projection axons from HVC to RA. (C) Identification of myelinated (orange) and unmyelinated axons in the HVC➔RA fiber tract. (D) DAB stain enters the axons at nodes of Ranvier (blue), but not at myelinated parts of the axon (orange), leaving myelinated segments unlabeled. (E) Comparison of diameters of descending projection axon and local collaterals of four HVC premotor neurons. (F) Diameters of labeled axons in EM images were measured along the shortest axis, minimizing systematic errors due to the unknown orientation of the fiber with respect to the imaging plane.

**Figure S3. Polychronous network sequences with HVC conduction delays and different initial conditions. Related to Figure 3 and Figure 5.** (A) Polychronous network sequence with HVC conduction delays and synchronously active starter neurons. (B) Zoom into the first 50 ms of the sequence in (A). (C) As in (A), starter neuron activity uniformly distributed within a 7 ms window. (D) Zoom into the first 50 ms of the sequence in (C). (E) As in (A), starter neurons are active at random time points within a 10 ms window. (F) Zoom into the first 50 ms of the sequence in (E). (G) Polychronous network sequence shown in Figure 3C (i.e., using HVC pathlength measurements and rodent conduction velocity); starter neurons are active synchronously. (H) Zoom into the first 50 ms of the sequence in (G).

## REFERENCES

Abeles, M. (1991). Corticonics: Neural Circuits of the Cerebral Cortex (Cambridge University Press).

Amador, A., Perl, Y.S., Mindlin, G.B., and Margoliash, D. (2013). Elemental gesture dynamics are encoded by song premotor cortical neurons. Nature 495, 59–64.

Amari, S. (1972). Learning patterns and pattern sequences by self-organizing nets of threshold elements. IEEE Trans Comp c-21, 1197–1206.

Andalman, A.S., Foerster, J.N., and Fee, M.S. (2011). Control of vocal and respiratory patterns in birdsong: dissection of forebrain and brainstem mechanisms using temperature. PLoS One 6, e25461.

Benezra, S.E., Narayanan, R.T., Egger, R., Oberlaender, M., and Long, M.A. (2018). Morphological characterization of HVC projection neurons in the zebra finch (Taeniopygia guttata). J Comp Neurol 526, 1673–1689.

Bienenstock, E. (1996). On the dimensionality of cortical graphs. J Physiol Paris 90, 251–256.

Bruno, R.M. (2011). Synchrony in sensation. Curr Opin Neurobiol 21, 701–708.

Bruno, R.M., and Sakmann, B. (2006). Cortex is driven by weak but synchronously active thalamocortical synapses. Science 312, 1622–1627.

Budd, J.M., Kovacs, K., Ferecsko, A.S., Buzas, P., Eysel, U.T., and Kisvarday, Z.F. (2010). Neocortical axon arbors trade-off material and conduction delay conservation. PLoS Comput Biol 6, e1000711.

Buzsaki, G., and Mizuseki, K. (2014). The log-dynamic brain: how skewed distributions affect network operations. Nat Rev Neurosci 15, 264–278.

Cannon, J., Kopell, N., Gardner, T., and Markowitz, J. (2015). Neural Sequence Generation Using Spatiotemporal Patterns of Inhibition. PLoS Comput Biol 11, e1004581.

Carr, C.E., and Konishi, M. (1988). Axonal delay lines for time measurement in the owl’s brainstem. Proc Natl Acad Sci U S A 85, 8311–8315.

Carr, C.E., and Konishi, M. (1990). A circuit for detection of interaural time differences in the brain stem of the barn owl. J Neurosci 10, 3227–3246.

Danish, H.H., Aronov, D., and Fee, M.S. (2017). Rhythmic syllable-related activity in a songbird motor thalamic nucleus necessary for learned vocalizations. PLoS One 12, e0169568.

Debrabant, K. (2010). Runge-Kutta methods for third order weak approximation of SDEs with multidimensional additive noise. Bit 50, 541–558.

Denk, W., Briggman, K.L., and Helmstaedter, M. (2012). Structural neurobiology: missing link to a mechanistic understanding of neural computation. Nat Rev Neurosci 13, 351–358.

Diesmann, M., Gewaltig, M.O., and Aertsen, A. (1999). Stable propagation of synchronous spiking in cortical neural networks. Nature 402, 529–533.

Fee, M.S., Kozhevnikov, A.A., and Hahnloser, R.H. (2004). Neural mechanisms of vocal sequence generation in the songbird. Ann N Y Acad Sci 1016, 153–170.

Fiete, I.R., Senn, W., Wang, C.Z., and Hahnloser, R.H. (2010). Spike-time-dependent plasticity and heterosynaptic competition organize networks to produce long scale-free sequences of neural activity. Neuron 65, 563–576.

Foster, D.J., and Wilson, M.A. (2007). Hippocampal theta sequences. Hippocampus 17, 1093–1099.

Galvis, D., Wu, W., Hyson, R.L., Johnson, F., and Bertram, R. (2018). Interhemispheric dominance switching in a neural network model for birdsong. J Neurophysiol 120, 1186–1197.

Gerstner, W., Kempter, R., van Hemmen, J.L., and Wagner, H. (1996). A neuronal learning rule for sub-millisecond temporal coding. Nature 383, 76–81.

Gibb, L., Gentner, T.Q., and Abarbanel, H.D. (2009). Inhibition and recurrent excitation in a computational model of sparse bursting in song nucleus HVC. J Neurophysiol 102, 1748–1762.

Goldman, M.S. (2009). Memory without feedback in a neural network. Neuron 61, 621–634.

Graber, M.H., Helmchen, F., and Hahnloser, R.H. (2013). Activity in a premotor cortical nucleus of zebra finches is locally organized and exhibits auditory selectivity in neurons but not in glia. PLoS One 8, e81177.

Hahnloser, R.H., Kozhevnikov, A.A., and Fee, M.S. (2002). An ultra-sparse code underlies the generation of neural sequences in a songbird. Nature 419, 65–70.

Hahnloser, R.H., Kozhevnikov, A.A., and Fee, M.S. (2006). Sleep-related neural activity in a premotor and a basal-ganglia pathway of the songbird. J Neurophysiol 96, 794–812.

Hamaguchi, K., Tanaka, M., and Mooney, R. (2016). A Distributed Recurrent Network Contributes to Temporally Precise Vocalizations. Neuron 91, 680–693.

Helmstaedter, M., Staiger, J.F., Sakmann, B., and Feldmeyer, D. (2008). Efficient recruitment of layer 2/3 interneurons by layer 4 input in single columns of rat somatosensory cortex. J Neurosci 28, 8273–8284.

Hirsch, J.A., and Gilbert, C.D. (1991). Synaptic physiology of horizontal connections in the cat’s visual cortex. J Neurosci 11, 1800–1809.

Hodgkin, A.L., and Huxley, A.F. (1952). A quantitative description of membrane current and its application to conduction and excitation in nerve. J Physiol 117, 500–544.

Hursh, J.B. (1939). Conduction velocity and diameter of nerve fibers. American Journal of Physiology 127, 131–139.

Innocenti, G.M., Lehmann, P., and Houzel, J.C. (1994). Computational structure of visual callosal axons. Eur J Neurosci 6, 918–935.

Itskov, V., Curto, C., Pastalkova, E., and Buzsaki, G. (2011). Cell assembly sequences arising from spike threshold adaptation keep track of time in the hippocampus. J Neurosci 31, 2828–2834.

Izhikevich, E.M. (2006). Polychronization: computation with spikes. Neural Comput 18, 245–282.

Jeffress, L.A. (1948). A place theory of sound localization. J Comp Physiol Psychol 41, 35–39.

Jin, D.Z., Ramazanoglu, F.M., and Seung, H.S. (2007). Intrinsic bursting enhances the robustness of a neural network model of sequence generation by avian brain area HVC. J Comput Neurosci 23, 283–299.

Jun, J.K., and Jin, D.Z. (2007). Development of neural circuitry for precise temporal sequences through spontaneous activity, axon remodeling, and synaptic plasticity. PLoS One 2, e723.

Katlowitz, K.A., Picardo, M.A., and Long, M.A. (2018). Stable Sequential Activity Underlying the Maintenance of a Precisely Executed Skilled Behavior. Neuron 98, 1133–1140 e1133.

Kleinfeld, D., and Sompolinsky, H. (1988). Associative neural network model for the generation of temporal patterns. Theory and application to central pattern generators. Biophys J 54, 1039–1051.

Kornfeld, J., Benezra, S.E., Narayanan, R.T., Svara, F., Egger, R., Oberlaender, M., Denk, W., and Long, M.A. (2017). EM connectomics reveals axonal target variation in a sequence-generating network. Elife 6.

Kosche, G., Vallentin, D., and Long, M.A. (2015). Interplay of inhibition and excitation shapes a premotor neural sequence. J Neurosci 35, 1217–1227.

Laje, R., and Buonomano, D.V. (2013). Robust timing and motor patterns by taming chaos in recurrent neural networks. Nat Neurosci 16, 925–933.

Liewald, D., Miller, R., Logothetis, N., Wagner, H.J., and Schuz, A. (2014). Distribution of axon diameters in cortical white matter: an electron-microscopic study on three human brains and a macaque. Biol Cybern 108, 541–557.

Lombardino, A.J., and Nottebohm, F. (2000). Age at deafening affects the stability of learned song in adult male zebra finches. J Neurosci 20, 5054–5064.

Long, M.A., and Fee, M.S. (2008). Using temperature to analyse temporal dynamics in the songbird motor pathway. Nature 456, 189–194.

Long, M.A., Jin, D.Z., and Fee, M.S. (2010). Support for a synaptic chain model of neuronal sequence generation. Nature 468, 394–399.

Lorteije, J.A., Rusu, S.I., Kushmerick, C., and Borst, J.G. (2009). Reliability and precision of the mouse calyx of Held synapse. J Neurosci 29, 13770–13784.

Luczak, A., McNaughton, B.L., and Harris, K.D. (2015). Packet-based communication in the cortex. Nat Rev Neurosci 16, 745–755.

Lynch, G.F., Okubo, T.S., Hanuschkin, A., Hahnloser, R.H., and Fee, M.S. (2016). Rhythmic Continuous-Time Coding in the Songbird Analog of Vocal Motor Cortex. Neuron 90, 877–892.

Magee, J.C., and Cook, E.P. (2000). Somatic EPSP amplitude is independent of synapse location in hippocampal pyramidal neurons. Nat Neurosci 3, 895–903.

Markowitz, J.E., Liberti, W.A., 3rd, Guitchounts, G., Velho, T., Lois, C., and Gardner, T.J. (2015). Mesoscopic patterns of neural activity support songbird cortical sequences. PLoS Biol 13, e1002158.

Mauk, M.D., and Buonomano, D.V. (2004). The neural basis of temporal processing. Annu Rev Neurosci 27, 307–340.

Mello, G.B., Soares, S., and Paton, J.J. (2015). A scalable population code for time in the striatum. Curr Biol 25, 1113–1122.

Miller, R. (1996). Axonal Conduction Time and Human Cerebral Laterality. A Psychobiological Theory. (Amsterdam, The Netherlands: Harwood Academic Publisher).

Miri, A., Daie, K., Arrenberg, A.B., Baier, H., Aksay, E., and Tank, D.W. (2011). Spatial gradients and multidimensional dynamics in a neural integrator circuit. Nat Neurosci 14, 1150–1159.

Mooney, R., and Prather, J.F. (2005). The HVC microcircuit: the synaptic basis for interactions between song motor and vocal plasticity pathways. J Neurosci 25, 1952–1964.

Narayanan, R.T., Egger, R., Johnson, A.S., Mansvelder, H.D., Sakmann, B., de Kock, C.P., and Oberlaender, M. (2015). Beyond Columnar Organization: Cell Type- and Target Layer-Specific Principles of Horizontal Axon Projection Patterns in Rat Vibrissal Cortex. Cereb Cortex 25, 4450–4468.

Nottebohm, F., Stokes, T.M., and Leonard, C.M. (1976). Central control of song in the canary, Serinus canarius. J Comp Neurol 165, 457–486.

Okubo, T.S., Mackevicius, E.L., Payne, H.L., Lynch, G.F., and Fee, M.S. (2015). Growth and splitting of neural sequences in songbird vocal development. Nature 528, 352–357.

Pastalkova, E., Itskov, V., Amarasingham, A., and Buzsaki, G. (2008). Internally generated cell assembly sequences in the rat hippocampus. Science 321, 1322–1327.

Peh, W.Y., Roberts, T.F., and Mooney, R. (2015). Imaging auditory representations of song and syllables in populations of sensorimotor neurons essential to vocal communication. J Neurosci 35, 5589–5605.

Pehlevan, C., Ali, F., and Olveczky, B.P. (2018). Flexibility in motor timing constrains the topology and dynamics of pattern generator circuits. Nat Commun 9, 977.

Peters, A.J., Chen, S.X., and Komiyama, T. (2014). Emergence of reproducible spatiotemporal activity during motor learning. Nature 510, 263–267.

Picardo, M.A., Merel, J., Katlowitz, K.A., Vallentin, D., Okobi, D.E., Benezra, S.E., Clary, R.C., Pnevmatikakis, E.A., Paninski, L., and Long, M.A. (2016). Population-Level Representation of a Temporal Sequence Underlying Song Production in the Zebra Finch. Neuron 90, 866–876.

Plaza, S.M., Scheffer, L.K., and Chklovskii, D.B. (2014). Toward large-scale connectome reconstructions. Curr Opin Neurobiol 25, 201–210.

Pnevmatikakis, E.A., Soudry, D., Gao, Y., Machado, T.A., Merel, J., Pfau, D., Reardon, T., Mu, Y., Lacefield, C., Yang, W., et al. (2016). Simultaneous Denoising, Deconvolution, and Demixing of Calcium Imaging Data. Neuron 89, 285–299.

Prut, Y., Vaadia, E., Bergman, H., Haalman, I., Slovin, H., and Abeles, M. (1998). Spatiotemporal structure of cortical activity: properties and behavioral relevance. J Neurophysiol 79, 2857–2874.

Rajan, K., Harvey, C.D., and Tank, D.W. (2016). Recurrent Network Models of Sequence Generation and Memory. Neuron 90, 128–142.

Rushton, W.A. (1951). A theory of the effects of fibre size in medullated nerve. J Physiol 115, 101–122.

Sabatini, B.L., and Regehr, W.G. (1996). Timing of neurotransmission at fast synapses in the mammalian brain. Nature 384, 170–172.

Sachidhanandam, S., Sreenivasan, V., Kyriakatos, A., Kremer, Y., and Petersen, C.C. (2013). Membrane potential correlates of sensory perception in mouse barrel cortex. Nat Neurosci 16, 1671–1677.

Salami, M., Itami, C., Tsumoto, T., and Kimura, F. (2003). Change of conduction velocity by regional myelination yields constant latency irrespective of distance between thalamus and cortex. Proc Natl Acad Sci U S A 100, 6174–6179.

Schmitt, L.I., Wimmer, R.D., Nakajima, M., Happ, M., Mofakham, S., and Halassa, M.M. (2017). Thalamic amplification of cortical connectivity sustains attentional control. Nature 545, 219–223.

Seung, H.S. (2012). Connectome: How the Brain’s Wiring Makes Us Who We Are (New York, NY: Houghton Mifflin Harcourt Publishing).

Shu, Y., Duque, A., Yu, Y., Haider, B., and McCormick, D.A. (2007). Properties of action-potential initiation in neocortical pyramidal cells: evidence from whole cell axon recordings. J Neurophysiol 97, 746–760.

Shu, Y., Hasenstaub, A., Duque, A., Yu, Y., and McCormick, D.A. (2006). Modulation of intracortical synaptic potentials by presynaptic somatic membrane potential. Nature 441, 761–765.

Sugihara, I., Lang, E.J., and Llinas, R. (1993). Uniform olivocerebellar conduction time underlies Purkinje cell complex spike synchronicity in the rat cerebellum. J Physiol 470, 243–271.

Swadlow, H.A. (1985). Physiological properties of individual cerebral axons studied in vivo for as long as one year. J Neurophysiol 54, 1346–1362.

Swadlow, H.A. (1994). Efferent neurons and suspected interneurons in motor cortex of the awake rabbit: axonal properties, sensory receptive fields, and subthreshold synaptic inputs. J Neurophysiol 71, 437–453.

Swadlow, H.A., Kocsis, J.D., and Waxman, S.G. (1980). Modulation of impulse conduction along the axonal tree. Annu Rev Biophys Bioeng 9, 143–179.

Swadlow, H.A., Waxman, S.G., and Weyand, T.G. (1981). Effects of variations in temperature on impulse conduction along nonmyelinated axons in the mammalian brain. Exp Neurol 71, 383–389.

Vu, E.T., Mazurek, M.E., and Kuo, Y.C. (1994). Identification of a forebrain motor programming network for the learned song of zebra finches. J Neurosci 14, 6924–6934.

Williams, S.R., and Stuart, G.J. (2002). Dependence of EPSP efficacy on synapse location in neocortical pyramidal neurons. Science 295, 1907–1910.

Yildiz, I.B., and Kiebel, S.J. (2011). A hierarchical neuronal model for generation and online recognition of birdsongs. PLoS Comput Biol 7, e1002303.

