## Supplementary figures and images for "Local axonal conduction delays underlie precise timing of a neural sequence"

### Supplemental Figure 1

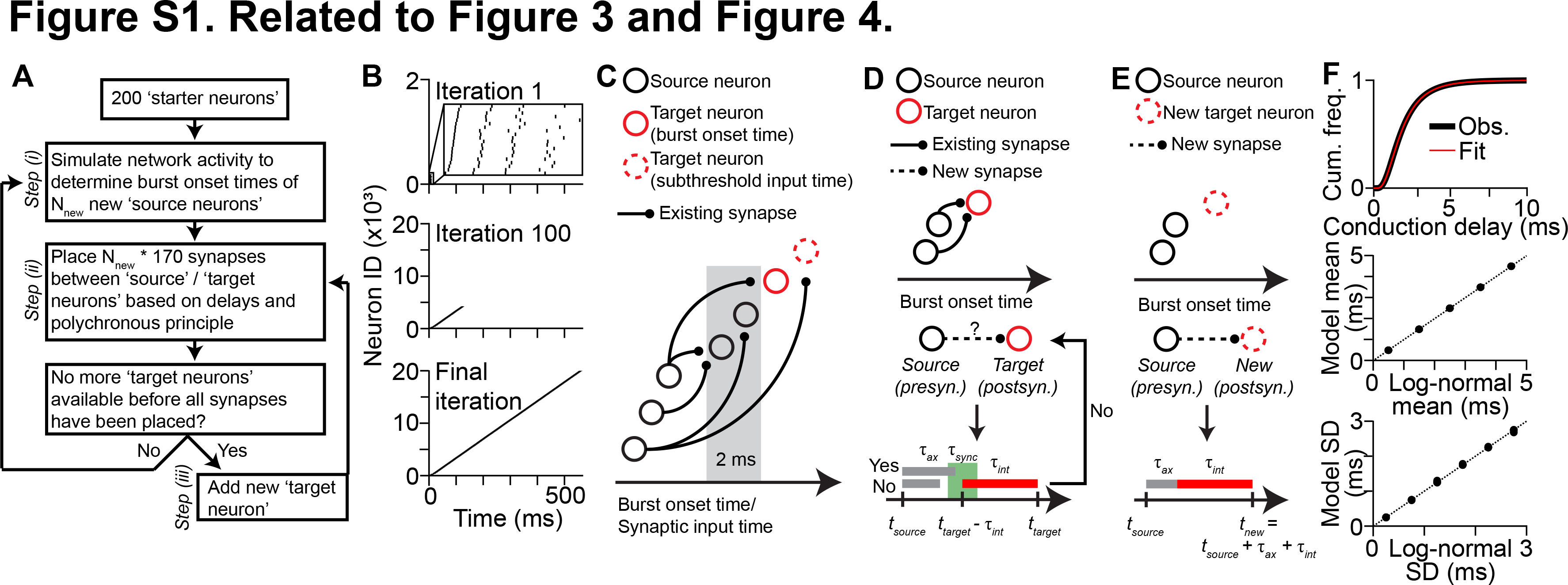

### Supplemental Figure 2

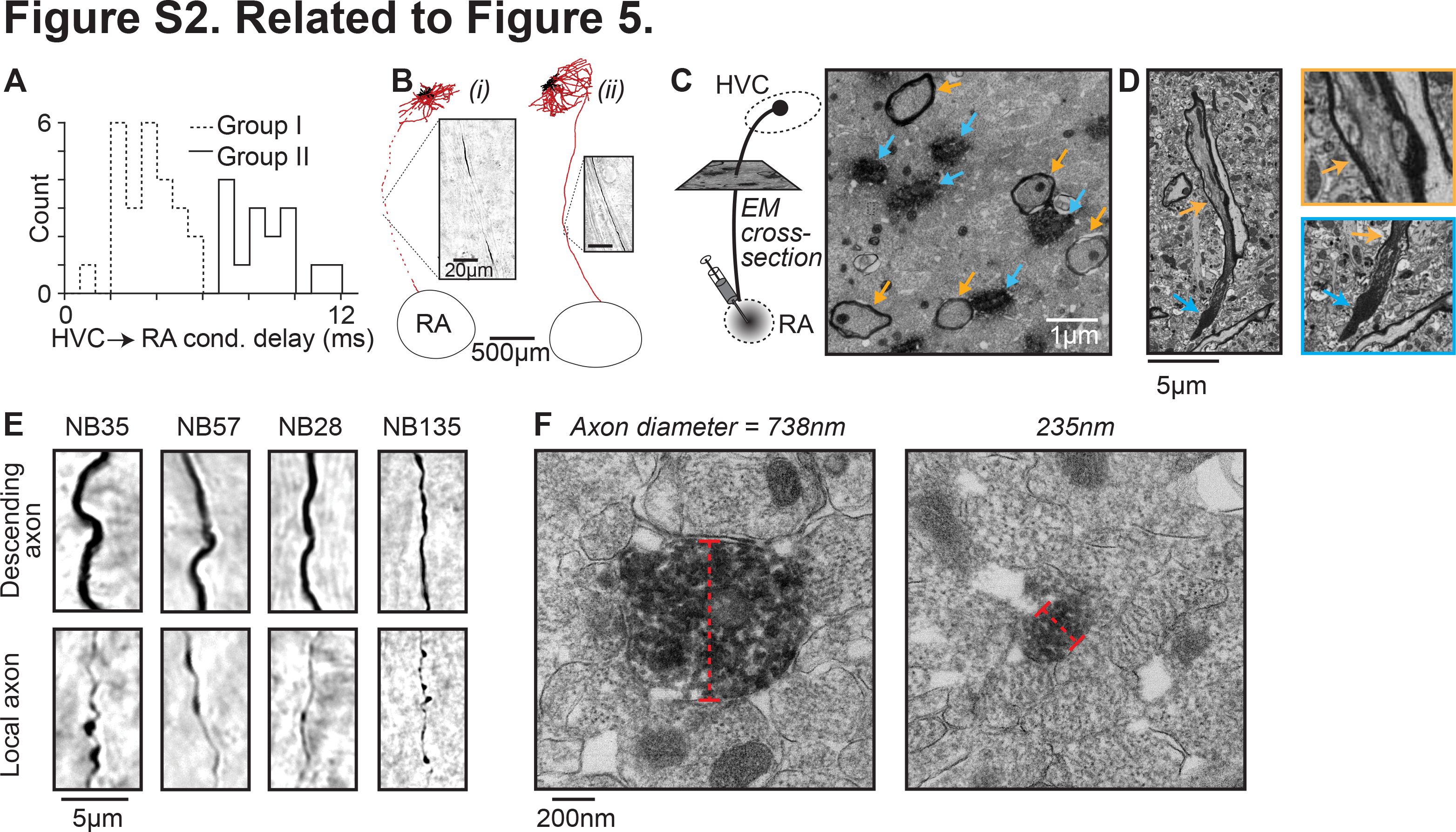

### Supplemental Figure 3

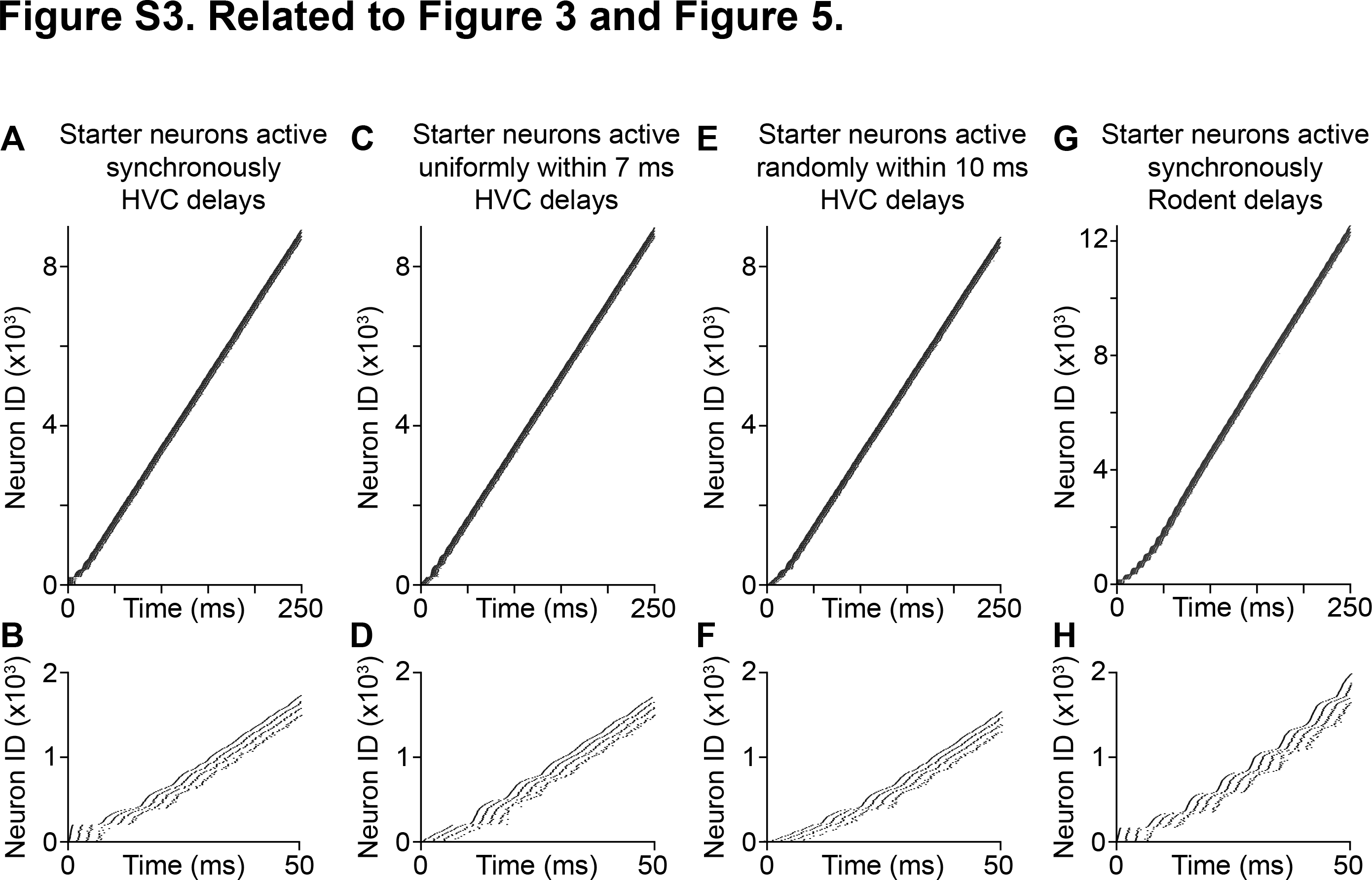
